# Elevated islet prohormone ratios as indicators of insulin dependency in islet transplant recipients

**DOI:** 10.1101/2021.10.17.464749

**Authors:** Yi-Chun Chen, Agnieszka M. Klimek-Abercrombie, Kathryn J. Potter, Lindsay P. Pallo, Galina Soukhatcheva, Dai Lei, Melena D. Bellin, C. Bruce Verchere

**Affiliations:** Department of Surgery, University of British Columbia, Vancouver, BC, Canada; BC Children’s Hospital Research Institute, Vancouver, BC, Canada; Novo Nordisk Canada Inc., Mississauga, Ontario, Canada; Montreal Clinical Research Institute, Montréal, Québec, Canada; Department of Pathology & Laboratory Medicine, University of British Columbia, Vancouver, BC, Canada; Department of Pediatrics and Surgery, University of Minnesota, Minneapolis, MN, USA; Centre of Molecular Medicine and Therapeutics, Vancouver, BC, Canada

## Abstract

Autologous pancreatic islet transplantation is an established therapy for patients with chronic pancreatitis. However, the long-term transplant outcomes are modest. Identifying indicators of graft function will aid the preservation of transplanted islets and glycemic control. To this end, we analyzed beta cell prohormone peptide levels in a retrospective cohort of total pancreatectomy autologous islet transplant patients (n=28). Proinsulin-to-C-peptide (PI/C) and proIAPP-to-total IAPP (proIAPP/IAPP) ratios measured at 3 months post-transplant were significantly higher in patients who remained insulin dependent at 1 year follow-up. In a mouse model of human islet transplantation, recipient mice that later became hyperglycemic displayed significantly higher PI/C ratios than mice that remained normoglycemic. Histological analysis of islet grafts showed reduced insulin- and proinsulin-positive area, but elevated glucagon-positive area in grafts that experienced greater secretory demand. Increased prohormone convertase 1/3 was detected in glucagon-positive cells, and glucagon-like peptide 1 (GLP-1) area was elevated in grafts from mice that displayed hyperglycemia or elevated plasma PI/C ratios, demonstrating intra-islet incretin production in metabolically challenged human islet grafts. These data indicate that in failing grafts, alpha cell prohormone processing is likely altered, and incomplete beta cell prohormone processing may be an early indicator of insulin dependency.

## Introduction

Pancreatic human islet transplant is an effective treatment for autoimmune-mediated diabetes or chronic pancreatitis^1,2^. After decades of refinements in surgical procedures, 50% of patients who received allo-transplant remained insulin independent 5 years post-transplant with modern immunosuppression protocols^3^, and around 30% of patients who underwent total pancreatectomy with islet autotransplantation (TPIAT) remained insulin independent 2 years post-surgery^1^. Other than insulin dependency, clinical assessments such as Igls, β-score^4,5^, hypoglycemic occurrence, homeostasis model assessment (HOMA) 2-B%, secretory units of islets in transplantation^6^, have been proposed to evaluate graft function. However, whether these indicators can be used to predict transplant outcomes remains to be tested. It has been reported that higher transplanted islet mass is associated with insulin independency^7^. Because TPIAT patients often received a sub-optimal islet mass, chronic metabolic stress on the beta cells is hypothesized to be one contributor to islet graft decline and lead to the return of insulin dependency. Metabolic assessments such as acute insulin response during intravenous glucose tolerance test at 3 months post-transplant or second-phase insulin secretion during intraperitoneal glucose tolerance test (IPGTT) have been reported to forecast long-term transplant outcomes^8^. The discovery of simpler and predictive indicators could benefit glycemic management in islet transplant patients and ultimately aid the preservation of graft function.

Beta-cell peptide prohormones are promising indicators of islet graft function and may be used to estimate transplant outcomes. Synthesized as larger precursor peptides, proinsulin and pro-islet amyloid polypeptide (IAPP) are processed by prohormone convertases and carboxypeptidase E to yield mature insulin, C-peptide, and IAPP. It has been shown that elevation of plasma proinsulin-to-C-peptide (PI/C) ratio and proIAPP-to-total IAPP (proIAPP/IAPP) ratio precede the development of both type 1 diabetes (T1D) and type 2 diabetes. ProIAPP/IAPP ratio was elevated in islet allotransplant patients compared to control immunosuppressant-treated individuals^9^. In patients who underwent allotransplantation and later developed hyperglycemia, higher plasma intact and split 32-33 proinsulin levels were detected^10^. Interestingly, plasma total proinsulin levels remained unchanged, and proinsulin-to-insulin ratios were lower in patients who remained insulin-independent^11^. Upon arginine stimulation, proinsulin processing efficiency is reduced in insulin-dependent allotransplant patients compared to insulin-independent patients^12^. These data suggest that impaired beta cell prohormone processing is associated with graft dysfunction. In this study, we tested whether PI/C or proIAPP/IAPP ratios can be used to evaluate islet graft function before the return of insulin dependency in a retrospective cohort of TPIAT patients.

Herein, we report elevated PI/C ratios in autologous islet transplant recipients who fail to achieve insulin independence at 1 year post-TPIAT. To gain insight into mechanisms contributing to islet graft dysfunction, we also performed optimal and sub-optimal human islet transplantation in immunodeficient mice made hyperglycemic by the beta-cell toxin streptozotocin (STZ) and measured human PI/C and proIAPP/IAPP ratios. We also performed immunohistological analysis on the islet grafts to evaluate prohormone processing patterns in islet cells. Our comprehensive study on prohormone processing machinery in a retrospective cohort of TPIAT patients and in an experimental setting could aid in the establishment of predictive biomarkers, and provide valuable information for the design of cell therapies to preserve graft function in islet transplantation.

## Material and methods

### Clinical human islet transplant

A cohort of 28 patients (age >11 years) undergoing total pancreatectomy with islet autotransplantation at the University of Minnesota from 2014-2015 were prospectively enrolled in a 1-year cohort study to islet engraftment^13^ At the time of TPIAT, all patients are started on insulin therapy, with insulin adjusted to target near euglycemia. After 3 months post-transplant, subcutaneous insulin is slowly weaned if tolerated, if patients maintain target blood glucoses of 80-125 mg/dL fasting, and <150-180 post-prandial, with a hemoglobin A1c ≤6.5%.. We utilized stored biospecimens from this study to retrospectively analyze the relationship between transplanted islet mass, pancreatic beta cell hormone levels (proinsulin, C-peptide, proIAPP_1-48_, and IAPP levels), metabolic outcomes (blood glucose, Hb_A1c_, glucose and C-peptide areas under the curve during a mixed meal tolerance test, acute C-peptide response to glucose and glucose potentiated acute C-peptide response to arginine during intravenous glucose tolerance test^13^), and insulin dependency at 90 days and one year post-transplant (Figure 1A). The study was approved by the University of Minnesota and the University of British Columbia institutional review boards. Informed consent or patient asset with parental informed consent was obtained.

**Figure 1.**
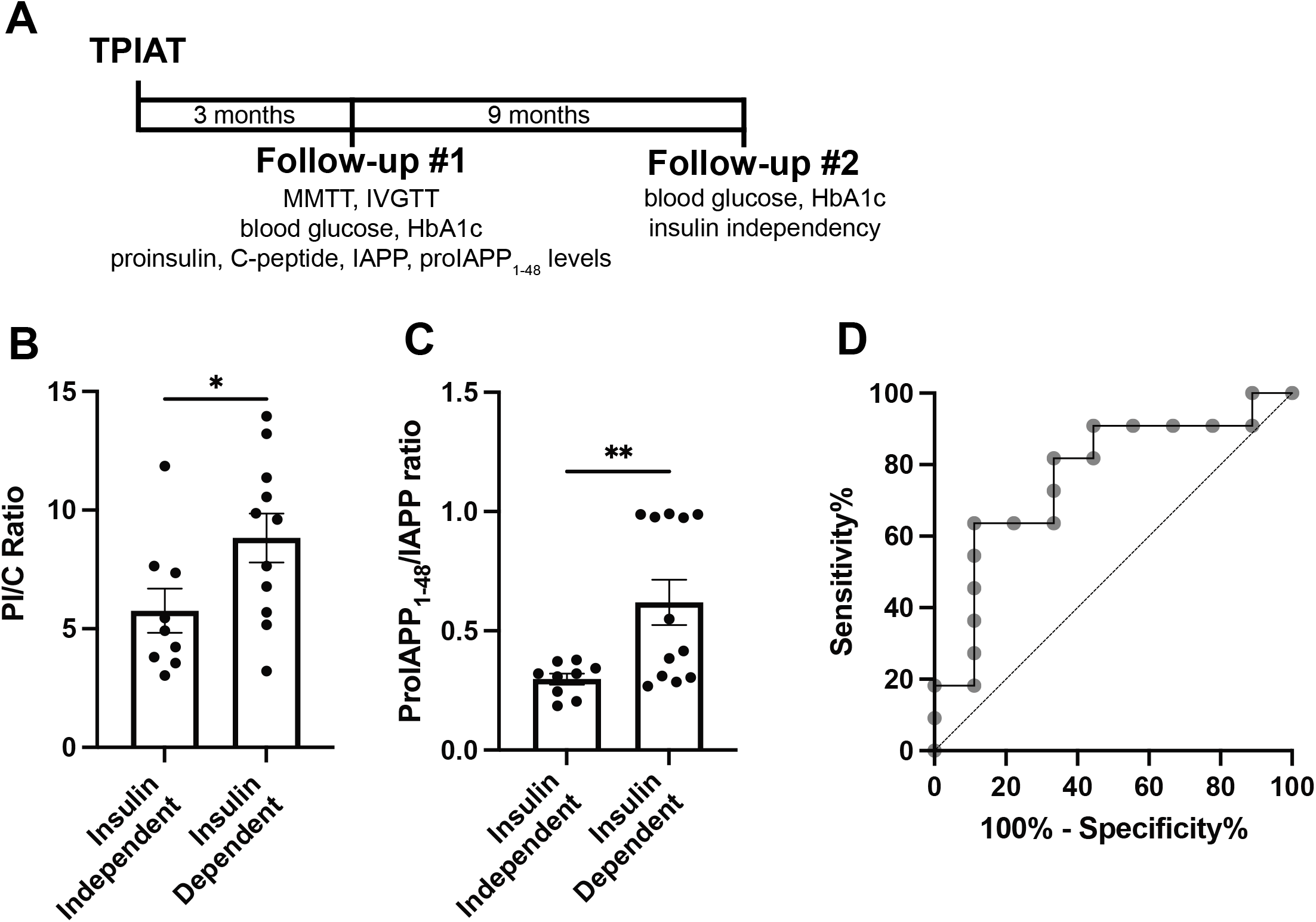
Islet prohormone-to-hormone ratios were elevated in TPIAT patients remained insulin-dependent. A) Timeline of a retrospective cohort of 28 patients undergoing total pancreatectomy with islet autotransplantation (TPIAT). B-C) Patients were separated into 2 groups based on insulin-dependency at 1 year post surgery, and proinsulin, C-peptide, proIAPP_1-48_, and amidated IAPP-like immunoreactivity were analyzed by ELISAs. Data were presented as mean ± SEM. *p < 0.05, **p < 0.01. D) Receiver-operating characteristic (ROC) analysis of plasma PI/C ratio and 1 year insulin-dependency of TPIAT patients.

### Optimal and sub-optimal human islet transplant in hyperglycemic and immunodeficient mice

Immunodeficient male NOD-*scid* IL2Rg^null^ (NSG) mice were purchased from The Jackson Laboratory (Bar Harbor, ME, USA) or BC Cancer Research Institute (Vancouver, BC, Canada) and maintained on a 14/10 h light/dark cycle, with a chow diet (6% fat, Teklab 2918, Huntingdon, UK) in BC Children’s Hospital Research Institute animal facility (Vancouver, BC, Canada). Prior to transplant, NSG mice were made diabetic with a single intraperitoneal injection of streptozotocin (STZ; Sigma, St Louis, MO, USA; 165-175 mg/kg body weight dissolved in pH4.5 citrate buffer. Mice received islet transplants 3 days following STZ injection if blood glucose levels ≥22 mM (28.6±0.4, n=51) as previously described. Unless otherwise noted, data presented are from mice that maintained normoglycemia for the 7 week duration of the experiment. Transplants were considered failure if blood glucose levels ≥ 11 mM during the 7 weeks post-transplant.

Human islets were obtained from University of Alberta Diabetes Institute IsletCore and Prodo Laboratories from a total of 7 donors (Supplementary Table 1). Human islets were allowed to recover overnight in 5.5 mM CMRL (Mediatech, Manassas, VA, USA) supplemented with 50 U/mL penicillin, 50mμg/mL streptomycin (Gibco/Fisher Scientific, Waltham, MA, USA), and 50 μg/mL gentamycin (Invitrogen, Waltham, MA, USA). Islets were hand-picked to >95% purity, counted, and transplanted under the kidney capsule of STZ-diabetic NSG mice as previously described^14^. Doses of either 250 (n=12), 300-330 (n=22, with 5 mice receiving 300 islets and 17 mice receiving 330 islets), 500 (n=10), or 750 (n=7) human islets from 7 donors were transplanted. For simplicity, the 300-330 islet group is hereafter referred to as 330 islets.

Mice that received human islet transplants were monitored weekly for body weight and fasting blood glucose levels, and fasting plasma were collected at 3- and 6-weeks post-transplant for beta-cell peptide hormone measurements. At 7-weeks post-transplant, a subset of mice received an intraperitoneal glucose tolerance test (n=26) using 1.5 g/kg body weight D-glucose (Sigma, St Louis, MO, USA). At the completion of the experiment (8 weeks post-transplant), upon excision of the graft-bearing kidney for histological analysis, all mice were allowed to recover and blood glucose levels were measured to ensure return of hyperglycemia.

### Circulating beta-cell peptide hormone measurements

Circulating human proinsulin and C-peptide levels in NSG mice were measured using Stellux Chemiluminescence ELISAs (Alpco, Salem, NH, USA). Human proinsulin levels in TPIAT patients were measured using TECO intact proinsulin ELISA (TECOmedical Group, Sissach, Switzerland), and human C-peptide levels in TPIAT patients were measured using an immuno-enzymometric assay on a TOSOH 2000 auto-analyzer (Tosoh Bioscience, Tokyo, Japan)^15^. Human proIAPP_1-48_ and amidated IAPP levels were measured using in-house developed Mesoscale Electroluminescence ELISAs (Rockville, MD, USA) as previously described^9^. For the patients with undetectable levels of amidated IAPP (4 out of 28 patients), the limit of detection of the assay divided by the square root of 2 was assigned to enable the inclusion of samples^16^.

### Immunohistological analysis

Islet grafts were excised at 7 weeks post-transplant, fixed in 4% paraformaldehyde-PBS solution (Sigma, St Louis, MO, USA), and paraffin-embedded. Sections (5 μm) were analyzed by immunofluorescence staining using primary antibodies against insulin, proinsulin, glucagon, synaptophysin, prohormone convertase 1/3 (PC1/3), and amidated glucagon-like peptide 1 (GLP-1), followed by highly cross-absorbed secondary Alexa Fluor-conjugated antibodies (Supplementary Table 2) as described previously^17^. Images were acquired using a Leica TCS SP5 confocal microscope (Leica Microsystems, Wetzlar, Germany). Once acquisition parameters were established using control and blindly selected experimental samples, 1 to 8 images were taken per islet graft, and 4 to 16 mice per experimental condition were analyzed. Antibody staining area and intensity were measured using automated CellProfiler pipelines^18^ in a blinded manner.

### Statistical analysis

The proinsulin/C-peptide ratios were calculated as proinsulin/C-peptide, and the proIAPP1-48/IAPP ratios were calculated as proIAPP_1-48_/(amidated IAPP+ proIAPP_1-48_). All data are presented as mean ± SEM. After validating Gaussian Distribution with Kolmogorov-Smirnov tests, a two-tailed t test with Welch’s correction, or Brown-Forsythe and Welch ANOVA tests were used. For correlation analysis, two-tailed Pearson correlation coefficients were calculated. For receiver-operating characteristic (ROC) analysis, area under the ROC curve and Youden’s index (specificity + sensitivity-1) were calculated to analyze discrimination of insulin dependency and to determine the optimal cut-off PI/C ratio. Data were expressed as mean ± SEM. ROC curve was reported as percentage. All analyses were performed using GraphPad Prism 9.

## Results

### Islet prohormone ratios as indicators of graft function and insulin dependency in TPIAT patients

We analyzed plasma PI/C and proIAPP/IAPP ratios in a cohort of 28 TPIAT patients (Table 1). Three months post-transplant, the PI/C ratio was negatively correlated with total islet equivalent (IEQ) and total islet number transplanted, and proIAPP/IAPP ratio was negatively correlated with total IEQ, total islet number transplanted, and total islet number transplanted per kg body weight (Table 2). These findings are in line with previous studies suggesting higher islet mass transplanted is associated with better graft function and insulin independency^7^. Because beta cell function is central to metabolic homeostasis, reduced graft function is often associated with poor metabolic outcome. Elevated PI/C ratio was associated with higher MMTT glucose area under the curve (AUC), and negatively associated with acute C-peptide response to glucose (ACRglu) and glucose-potentiated acute C-peptide response to arginine (ACRpot). Elevated proIAPP/IAPP ratio was negatively associated with ACRpot.

**Table 1:** Patient characteristics

|  |  |
| --- | --- |
| Age, mean (SD) | 35.5 (17.26) |
| Sex, n (%) |  |
| Female | 20/28 (71.42) |
| Male | 8/28 (28.57) |
| Baseline BMI, kg/m<sup>2</sup>, mean (SD) | 26.3 (6.30) |
| Baseline glycemic status, blood glucose levels (mg/dL), mean (SD) | 99.0 (17.44) |
| Total IEQ transplanted, mean (SD) | 358290.0 (175189.72) |
| IEQ per kg transplanted, mean (SD) | 5016.4 (2669.56) |
| Total islet number transplanted, mean (SD) | 389869.1 (182207.27) |
| Total islet number per kg, mean (SD) | 5601.1 (3722.34) |
| 2<sup>nd</sup> transplant site (peritoneal, mesentery, or subperitoneal), n (%) | 8/28 (28.57) |

**Table 2:** Correlation between PI/C and proIAPP/IAPP verses clinical variables

| *Clinical variable* | **PI/C** |  |  | **proIAPP<sub>1-48</sub>/IAPP** |  |  |
| --- | --- | --- | --- | --- | --- | --- |
|  | *Pearson r* | *R squared* | *P value* | *Pearson r* | *R squared* | *P value* |
| Weight (kg) | -0.3154 | 0.0995 | 0.1756 | -0.0032 | 1.08e-5 | 0.9887 |
| Height (cm) | -0.1723 | 0.0297 | 0.4675 | 0.2136 | 0.0456 | 0.3526 |
| Total IEQ | -0.5146 | 0.2649 | **0.0202** | -0.5706 | 0.3255 | **0.0069** |
| IEQ per kg | -0.3161 | 0.0999 | 0.1746 | -0.5293 | 0.2801 | **0.0136** |
| Total islet number | -0.3409 | 0.1162 | 0.1413 | -0.4486 | 0.2013 | **0.0414** |
| Insulin per kg | -0.0228 | 0.0005 | 0.9239 | -0.3374 | 0.1138 | 0.1347 |
| HbA1c day 90 | 0.3338 | 0.1114 | 0.1503 | 0.6394 | 0.4088 | **0.0018** |
| Insulin usage day 90 | 0.4535 | 0.2057 | **0.0446** | 0.4071 | 0.1657 | 0.0670 |
| Weight day 90 (kg) | -0.2899 | 0.0840 | 0.2151 | -0.0604 | 0.0036 | 0.7948 |
| Insulin usage day 90 per kg | 0.5963 | 0.3555 | **0.0055** | 0.5359 | 0.2872 | **0.0123** |
| Insulin Use 1 year | 0.4395 | 0.1932 | 0.0525 | 0.4199 | 0.1763 | 0.0653 |
| Weight 1 year | -0.2865 | 0.0821 | 0.2207 | -0.1527 | 0.0233 | 0.5203 |
| HbA1c 1 year | 0.3763 | 0.1416 | 0.1020 | 0.5196 | 0.2700 | **0.0189** |
| Age at TPIAT | -0.6051 | 0.3661 | **0.0047** | -0.3227 | 0.1041 | 0.1536 |
| Intra-portal IEQ | -0.4391 | 0.1928 | 0.0528 | -0.4255 | 0.1811 | 0.0545 |
| Intra-portal IEQ per kg | -0.3250 | 0.1056 | 0.1620 | -0.4360 | 0.1901 | **0.0482** |
| AUCglu | 0.7140 | 0.5098 | **0.0006** | 0.4217 | 0.1779 | 0.0721 |
| AUCcp | -0.2389 | 0.0571 | 0.3246 | -0.3951 | 0.1561 | 0.0941 |
| ACRglu | -0.5970 | 0.2885 | **0.0055** | -0.3554 | 0.1263 | 0.1138 |
| ACRpot | -0.5645 | 0.3187 | **0.0147** | -0.5288 | 0.2796 | **0.0199** |

Patients who remained insulin dependent 1-year post-surgery already displayed significantly higher PI/C ratios at 3 months post-surgery (Figure 1B and 1C). ROC analysis suggested that 3 months PI/C ratio is an acceptable indicator of 1 year insulin dependency (area under ROC curve = 0.78). Further analysis suggested a cut-off value of 1.847 to discriminate between insulin dependency and insulin independency at 1-year post-transplant (with Youden’s index of 2.5) (Figure 1D).

### PI/C ratio as a predictor of human islet transplant failure in mice

To model islet secretory stress and elucidate the cellular mechanism leading to elevated PI/C ratio, we transplanted groups of 250, 330, 500, and 750 hand-picked human islets from 7 donors (islet donor information listed in Supplementary Table 1) on the kidney capsule of immunodeficient mice made hyperglycemic with the beta-cell toxin STZ. We showed that mice that received fewer human islets displayed higher fasting blood glucose levels throughout the 7 week post-transplant timespan, and hyperglycemia in all mice resumed when the graft-bearing kidney was removed (Figure 2A). Although mice that received fewer human islets showed elevated glucose excursion during an i.p. glucose tolerance test. All mice were able to clear blood glucose swiftly, suggesting that transplanted human islets likely release insulin in a regulated manner in all groups (Figure 2B). Interestingly, at 3 weeks post-transplant, fasting plasma human proinsulin levels were significantly higher in mice that received 250 or 330 human islets compared to mice that received 750 human islets (Figure 2C), whereas human C-peptide levels were similar in all groups (Figure 2D). PI/C ratios were also significantly higher in mice that received 250, 330, or 500 human islets, compared to mice that received 750 human islets (Figure 2E), suggesting insufficient prohormone processing associated with lower islet transplant dose. Most importantly, mice that later become hyperglycemic (fasting blood glucose ≥11 mM during weeks 4-7 post-transplant) displayed significantly higher plasma human proinsulin level and PI/C ratios at 3 weeks post-transplant (Figure 2F and 2G). Both plasma human proinsulin and PI/C ratios were positively correlated with fasting blood glucose levels (Figure 2H and 2I). At 6 weeks post-transplant, plasma proinsulin level and PI/C ratios remained high in mice that received 250 or 330 human islets (Figure 2J and 2K).

**Figure 2.**
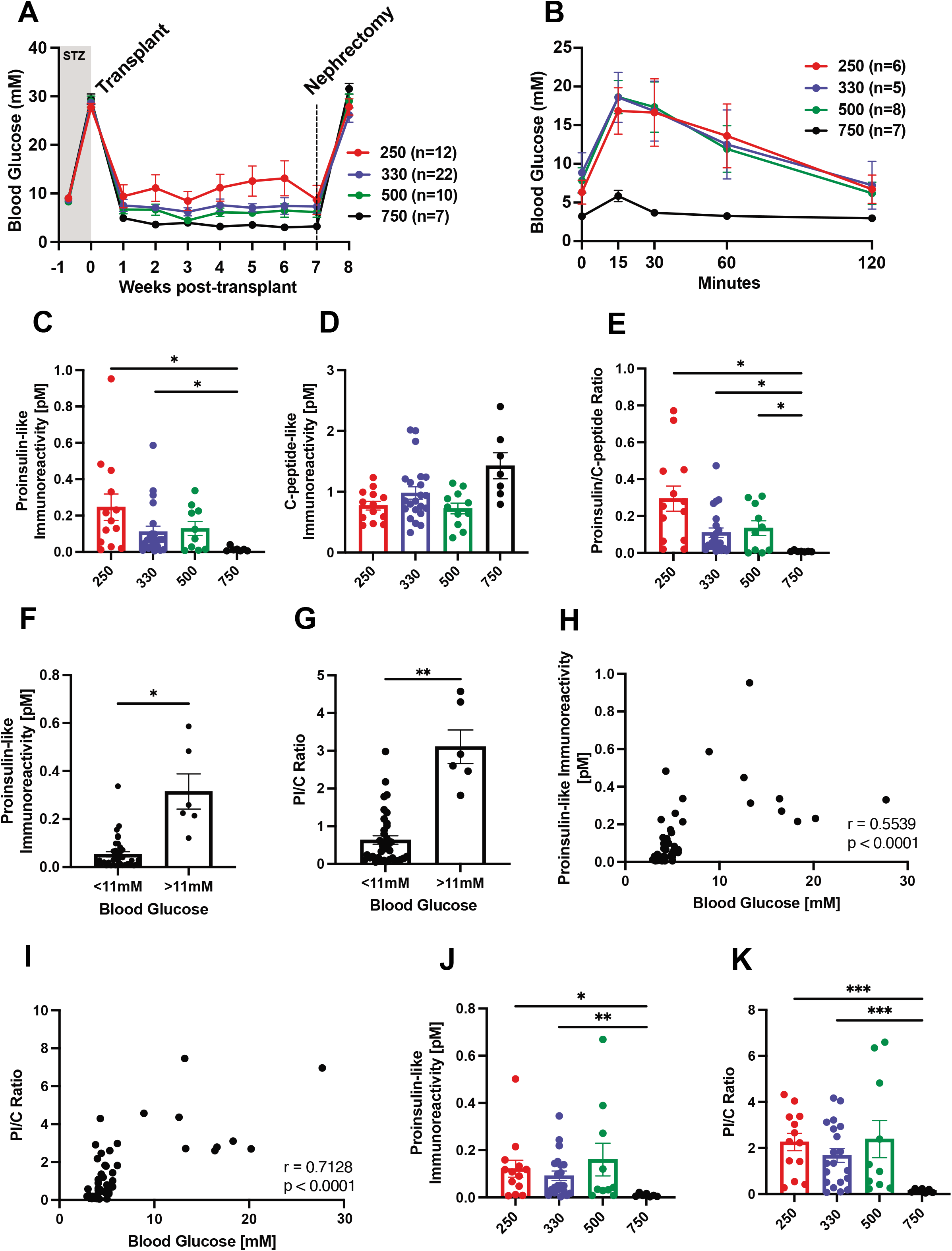
PI/C ratios were elevated in human islet transplanted mice later became hyperglycemic. A) NSG mice made hyperglycemic with streptozotocin (STZ) injection were transplanted with 250 (n=12), 330 (n=22), 500 (n=8), or 750 (n=7) hand-picked human islets. Body weights and fasting plasma glucose levels were measured weekly. B) Intraperitoneal glucose tolerance test was performed 7 weeks post-transplantation. Fasting plasma was collected at 3 weeks post-transplant for the measurements of C) human proinsulin, D) human C-peptide levels, and E) PI/C ratios. Mice were separated into two groups based on their transplant outcomes (fasting plasma glucose levels < 11mM or ≥ 11mM) between 4-7 weeks post-transplant, and F) fasting plasma human proinsulin levels and G) PI/C ratios at 3 weeks were compared. Correlation between fasting blood glucose and H) plasma human proinsulin levels or I) PI/C ratios in human islet transplanted mice at 3 weeks post-transplant. Fasting plasma was collected at 6 weeks post-transplant for the measurements of J) human proinsulin, K) PI/C ratios. Data were presented as mean ± SEM. *p < 0.05, **p < 0.01.

### Prohormone ratios associated with transplanted human islet dose in diabetic mice

ProIAPP is processed in parallel to proinsulin along the secretory pathway^19^ and elevated proIAPP/IAPP ratios have been used as an indicators of beta-cell dysfunction^9^. We measured an intermediate form of proIAPP, proIAPP_1-48_, and mature amidated IAPP levels using in-house human-(pro)IAPP specific ELISAs. Similar to proinsulin, we showed that while mature and amidated IAPP levels were comparable in all groups, proIAPP_1-48_ levels and the proIAPP/IAPP ratios were elevated in mice transplanted with 250 or 330 human islets (Figure 3A-3C). As predicted, plasma human proinsulin and proIAPP levels are positively correlated, and PI/C and proIAPP/IAPP ratios are also correlated (Figure 3D and 3E).

**Figure 3.**
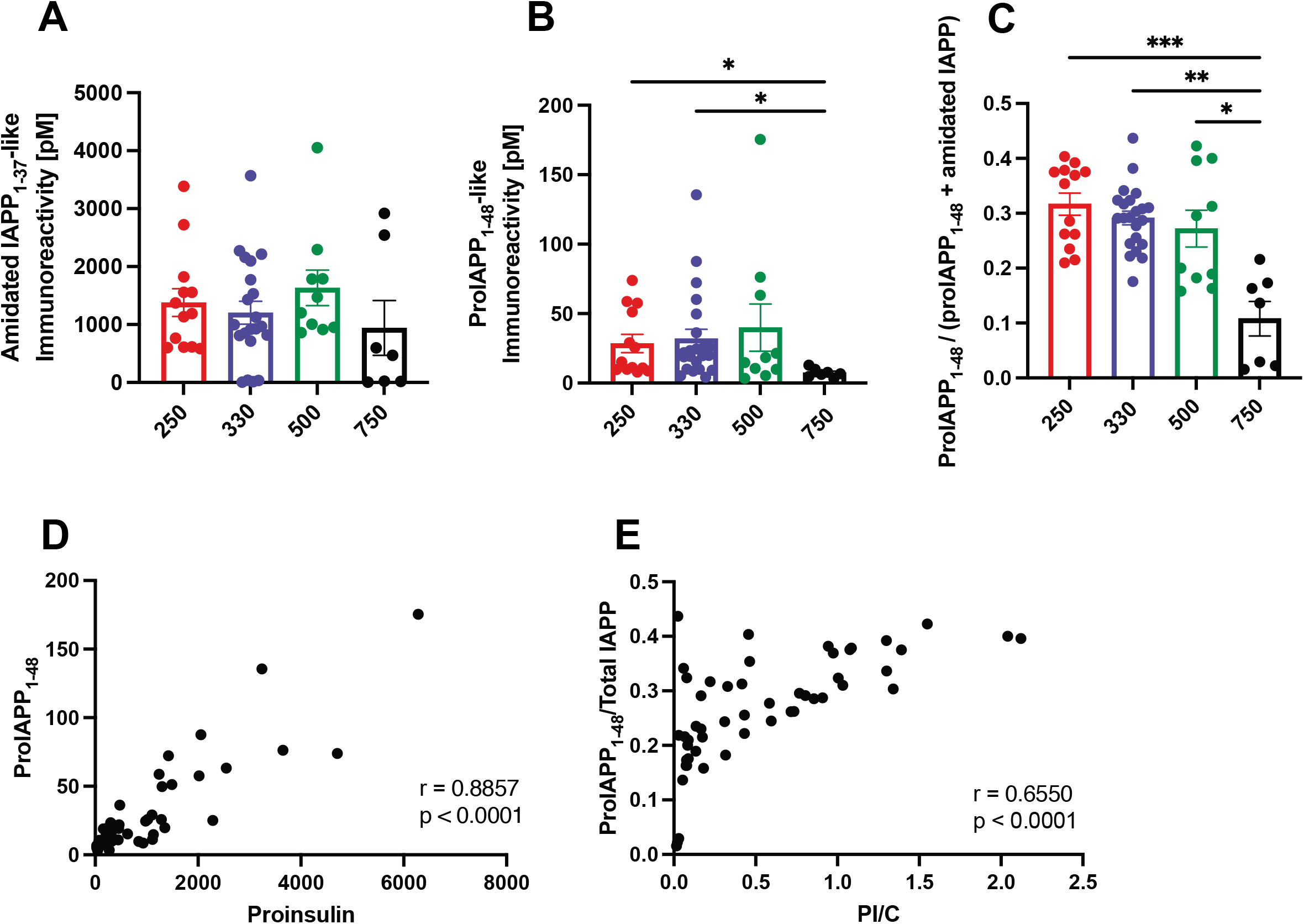
proIAPP/IAPP ratios were elevated in mice that received fewer human islets. Fasting plasma from mice that received human islet transplants was collected for the measurements of human A) amidated IAPP, B) proIAPP_1-48_ immunoreactivity, and C) proIAPP_1-48_/(amidated IAPP + proIAPP_1-48_) ratios at 6 weeks post-transplant. Correlations between D) proinsulin and proIAPP_1-48_, and E) PI/C and proIAPP/IAPP measured at 6 weeks post-transplant were analyzed. Data were presented as mean ± SEM. *p < 0.05, **p < 0.01.

### Sub-optimal human islet grafts have increased glucagon expression

To analyze islet cell composition in grafts experiencing higher secretory demand, we performed immunohistological analysis to examine the morphology of recovered islet grafts (Figure 4A). We showed that while insulin-staining positive (INS^+^) area and insulin staining intensity in islet (marked by synaptophysin-staining, SYT^+^) were not significantly reduced (area: p= 0.067, intensity: p= 0.11) (Figure 4B and 4C), glucagon-positive (GCG^+^) area was significantly elevated in islet grafts from mice that received less than 750 islets (Figure 4D), and glucagon staining intensity was significantly increased in mice that displayed hyperglycemia or elevated PI/C ratios (Figure 5E).

**Figure 4.**
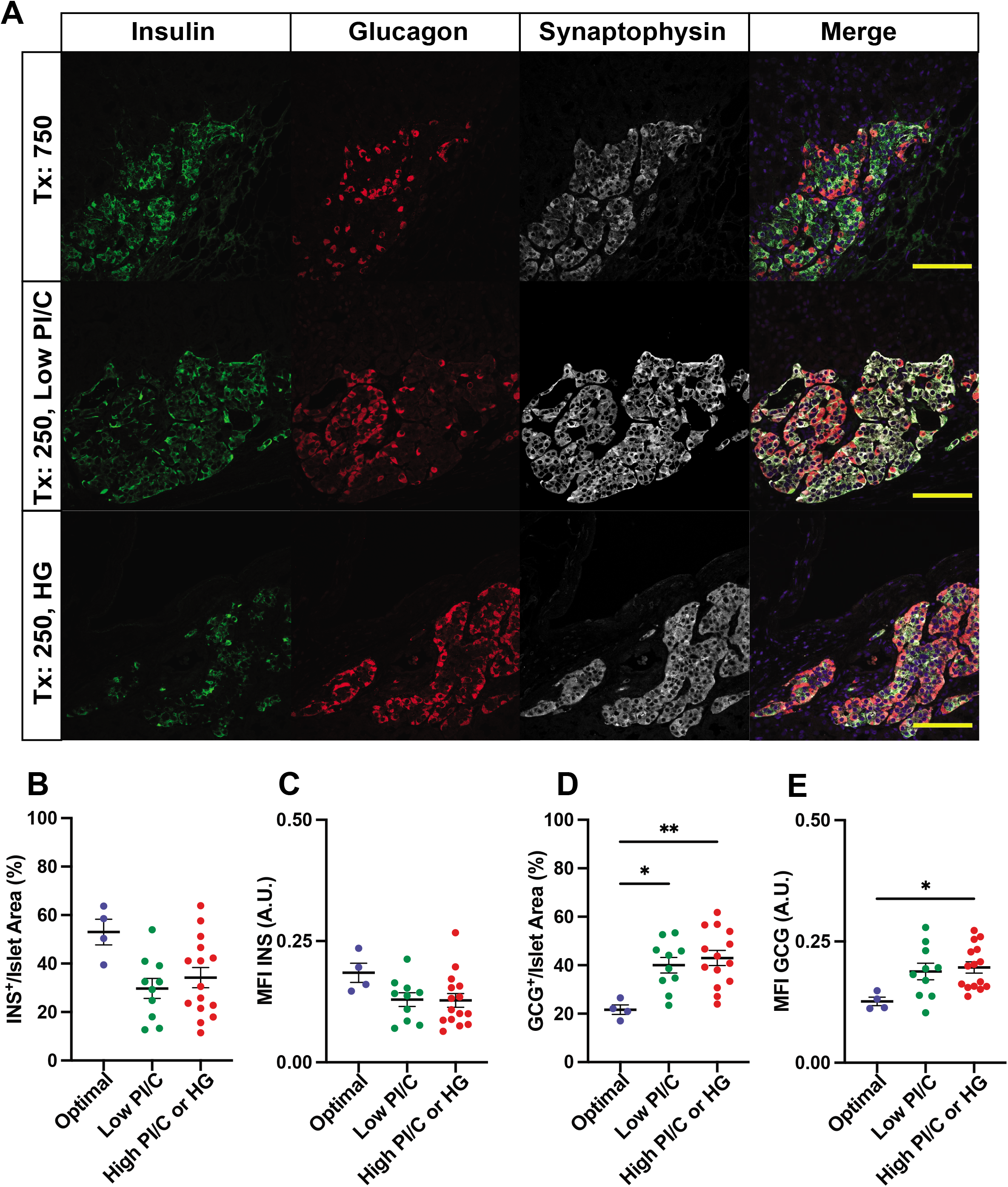
Glucagon-positive cell area in islet graft is elevated in human islet grafts from mice that displayed high PI/C ratios or hyperglycemia. A) Islet grafts were recovered by nephrectomy at 7 weeks post-transplant, and analyzed by immunofluorescence staining using antibodies against insulin, glucagon, and synaptophysin. B) Insulin-positive (INS^+^) area over synaptophysin-positive islet area, C) insulin staining intensity, D) glucagon-positive (GCG^+^) area over synaptophysin-positive islet area, and E) glucagon staining intensity were analyzed. Each data point represents the average of 1-7 images taken from a kidney-bearing islet graft from one mouse. Scale bar = 10μm Data were presented as mean ± SEM. *p < 0.05, **p < 0.01.

**Figure 5.**
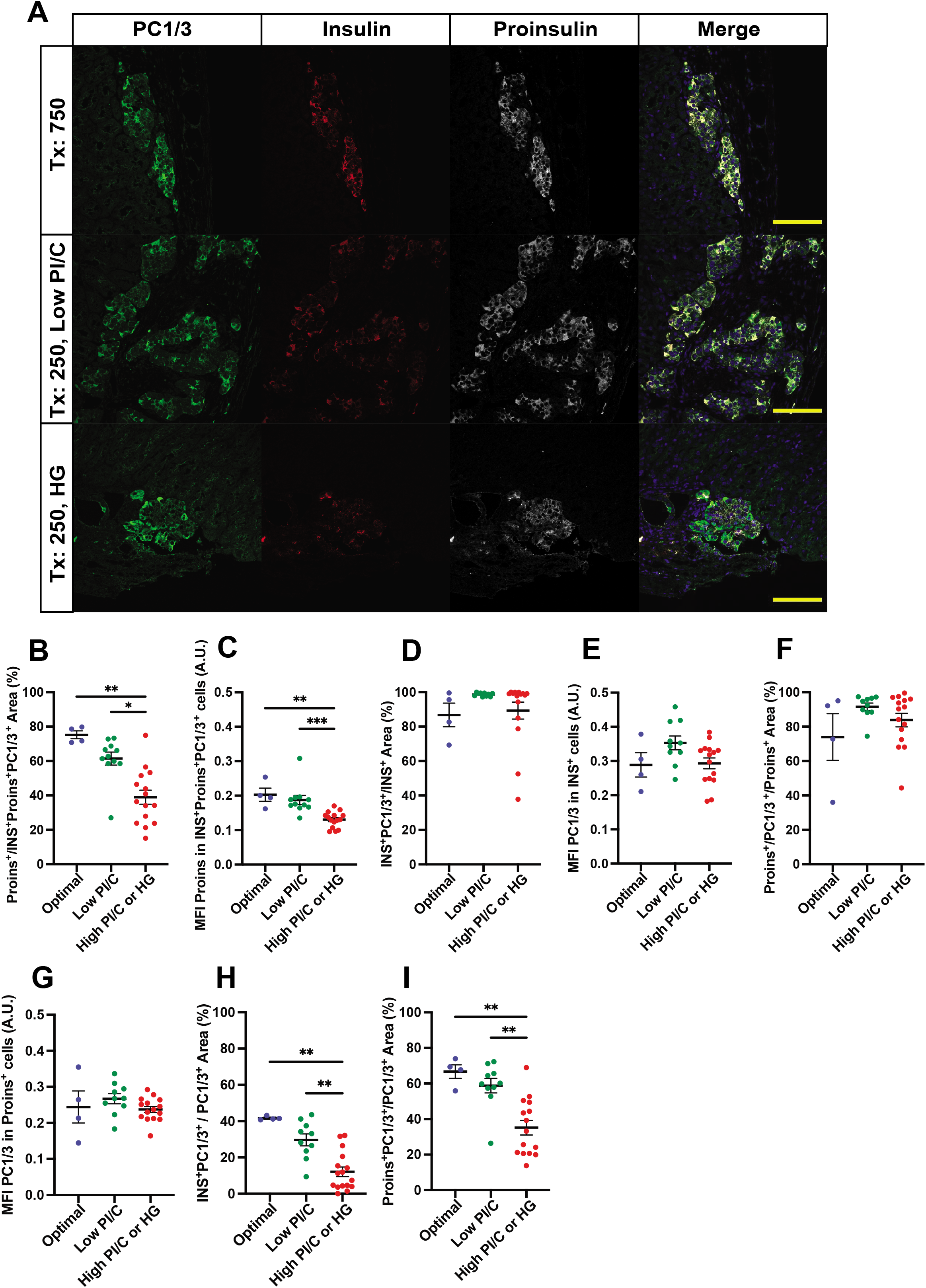
PC1/3 area or staining intensity was not reduced in insulin-positive or proinsulin-positive cells in human islet grafts from mice that displayed high PI/C ratios or hyperglycemia. A) Islet grafts were analyzed by immunofluorescence staining using antibodies against PC1/3, insulin, and proinsulin. B) Proinsulin-positive area over islet area (the combination insulin, proinsulin, and PC1/3-staining positive area), and C) proinsulin staining intensity were analyzed. PC1/3 area and intensity in insulin-positive cells D-E) and proinsulin-positive cells F-G) were analyzed. Proportional I) PC1/3 and insulin double-positive cells area and J) PC1/3 and proinsulin double-positive cells area in PC1/3-positive area were analyzed. Scale bar = 10μm Data were presented as mean ± SEM. *p < 0.05, **p < 0.01.

### Impaired prohormone processing in sub-optimal human islet grafts

Our observation of elevated plasma human PI/C and proIAPP/IAPP ratios in mice that received fewer human islets hints toward inefficient prohormone processing. To test this, we performed immunostaining of proinsulin, insulin, and PC1/3 in recovered islet grafts (Figure 5A). Unexpectedly, our data showed that proinsulin-staining positive (Proins^+^) area over islet area (INS^+^Proins^+^PC1/3^+^) was reduced in mice that displayed hyperglycemia or elevated PI/C ratios (Figure 5B). Similarly, proinsulin staining intensity was also reduced in mice that displayed hyperglycemia or elevated PI/C ratios (Figure 5C). One plausible explanation is that absolute proinsulin content in islets under secretory stress in mice that received fewer islets is indeed reduced, but these stressed islets were able to release unprocessed or partially processed insulin into the bloodstream.

In order to analyze proinsulin processing efficiency in transplanted human islet beta cells, we examined PC1/3 expression patterns in INS^+^ and Proins^+^ cells. We found that the percent PC1/3-staining positive (PC1/3^+^) area and PC1/3 intensity in INS^+^ cells were comparable in all groups (Figure 5D and 5E). Percent PC1/3^+^ area and intensity in Proins^+^ cells also showed similar patterns (Figure 5F and 5G). Our data suggest that prohormone processing enzyme PC1/3 expression levels are not changed in beta cells of human islets under secretory stress in vivo. Interestingly, the distribution of PC1/3^+^ cells in islet cell types was altered. We found that the INS^+^PC1/3^+^ area as a percent of PC1/3^+^ area is reduced in islets of mice that received fewer human islets (Figure 5H). Likewise, the Proins^+^PC1/3^+^ area as a proportion of PC1/3^+^ area is also reduced (Figure 5I).

### Elevated blood glucose and PI/C ratios in murine recipients sub-optimal human islet transplants

To analyze PC1/3 expression in alpha cells, we performed PC1/3 and glucagon double staining. Because proglucagon is processed by PC1/3 to GLP-1, we also analyzed GLP-1 expression pattern (using an antibody targeting amidated mature GLP-1) as a surrogate readout of PC1/3 activity in alpha cells. We found that PC1/3 is expressed in alpha cells, albeit at a lower level compared to beta cells (Figure 6A). When comparing alpha cells in islet grafts of mice that received 750 compared to those that received fewer than 750 human islets, we observed a non-significant (p= 0.16) increase in the GCG^+^PC1/3^+^ area as a percent of PC1/3^+^ area (Figure 6B). This observation complements our data in Figure 6, showing an increased proportion of PC1/3^+^ alpha cells. We found that the PC1/3^+^ area as a proportion of GCG^+^ area was not changed in human islet grafts in mice that became hyperglycemic or displayed elevated PI/C ratios (Figure 6C), and intensity of PC1/3 staining in GCG^+^ cells were not different from mice that received 750 human islets grafts (Figure 6D). GLP-1-positive (GLP-1^+^) area (as a percent of GCG^+^ area was tended to be higher in mice that developed hyperglycemia or had elevated PI/C ratios, but did not reach statistical significance (p= 0.14) (Figure 6E). GLP-1 staining intensity in GCG^+^ cells was slightly higher in hyperglycemic mice or mice that had higher PI/C ratios compared to those with normal PI/C ratios (Figure 7F). When we assessed GLP-1 staining in alpha cells by examining GLP-1^+^ cells in INS^-^ cells (Figure 7A), we observed that GLP-1 staining intensity was higher in INS^-^ cells of human islet grafts from hyperglycemic mice or mice that displayed higher plasma PI/C ratios (Figure 7B). It is likely that a proportion of alpha cells in the sub-optimal islet transplant group increase PC1/3 expression and generate GLP-1, likely explaining the reduced GCG staining intensity in PC1/3^+^GCG^+^ cells and GLP-1^+^GCG^+^ cells (Figure 6G and 6H). When analyzing GLP-1^+^ area in islet grafts by calculating GLP-1^+^ area as a percent of INS^+^ and GCG^+^ area, we found that islet GLP-1^+^ area was significantly higher in hyperglycemic mice or mice that had higher plasma PI/C ratios (Figure 7C). GLP-1^+^INS^-^ area in islet grafts was also elevated in mice that displayed hyperglycemia or higher PI/C ratios (Figure 7D).

**Figure 6.**
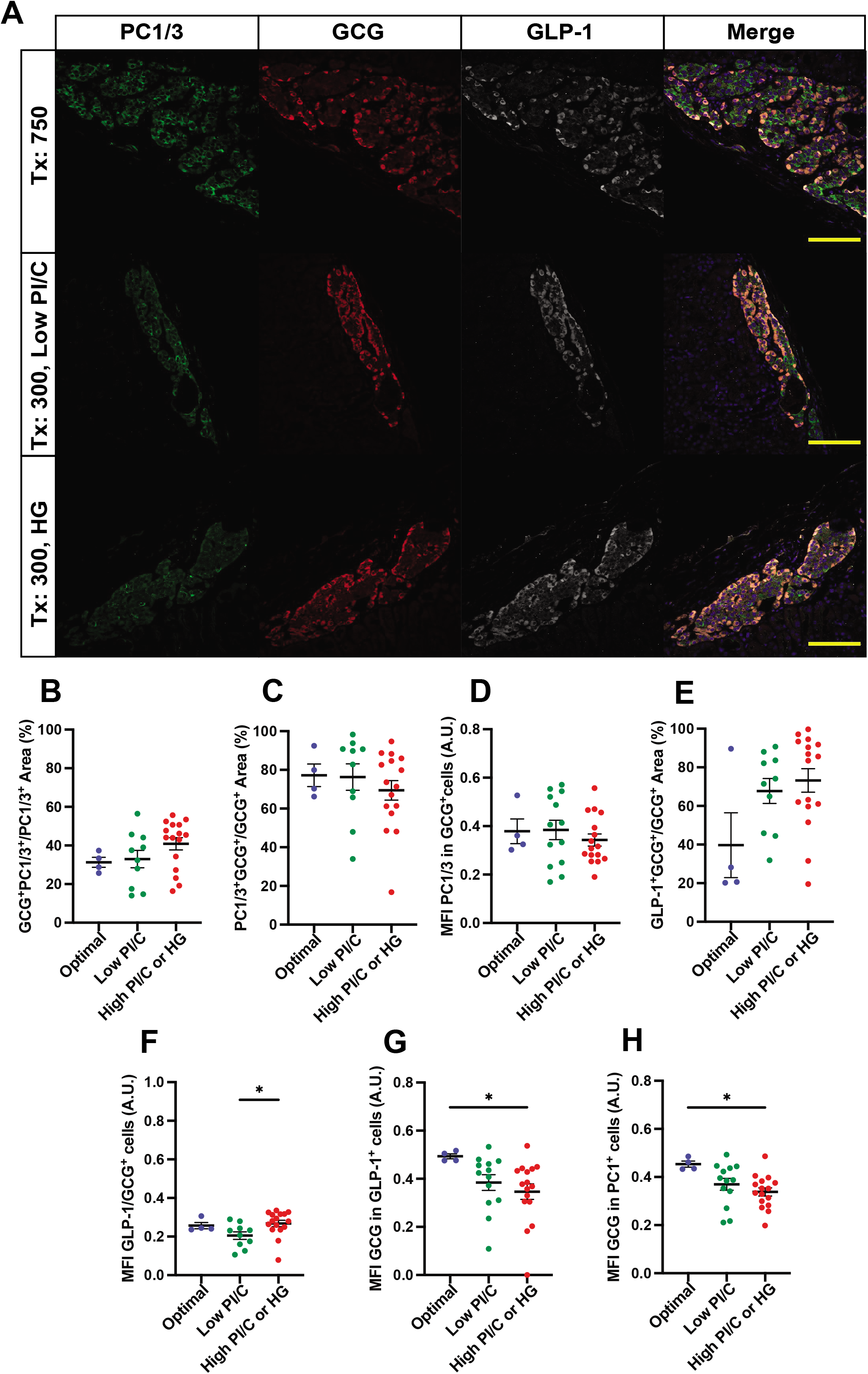
PC1/3 area or staining intensity was not changed in glucagon-positive cells in human islet grafts from mice that displayed high PI/C ratios or hyperglycemia. A) Islet grafts were analyzed by immunofluorescence staining using antibodies against PC1/3, glucagon, and amidated GLP-1. Proportional PC1/3 and glucagon double-positive area in B) PC1/3-positive area, and C) glucagon-positive area were analyzed. D) PC1/3 staining intensity in glucagon-positive cells were analyzed. E) Proportional GLP-1 and glucagon double-positive area in glucagon-positive area were analyzed. F) GLP-1 staining intensity in glucagon-positive cells, glucagon staining intensity in G) GLP-1-positive cells and H) PC1/3-positive cells were analyzed. Scale bar = 10μm Data were presented as mean ± SEM. *p < 0.05, **p < 0.01.

**Figure 7.**
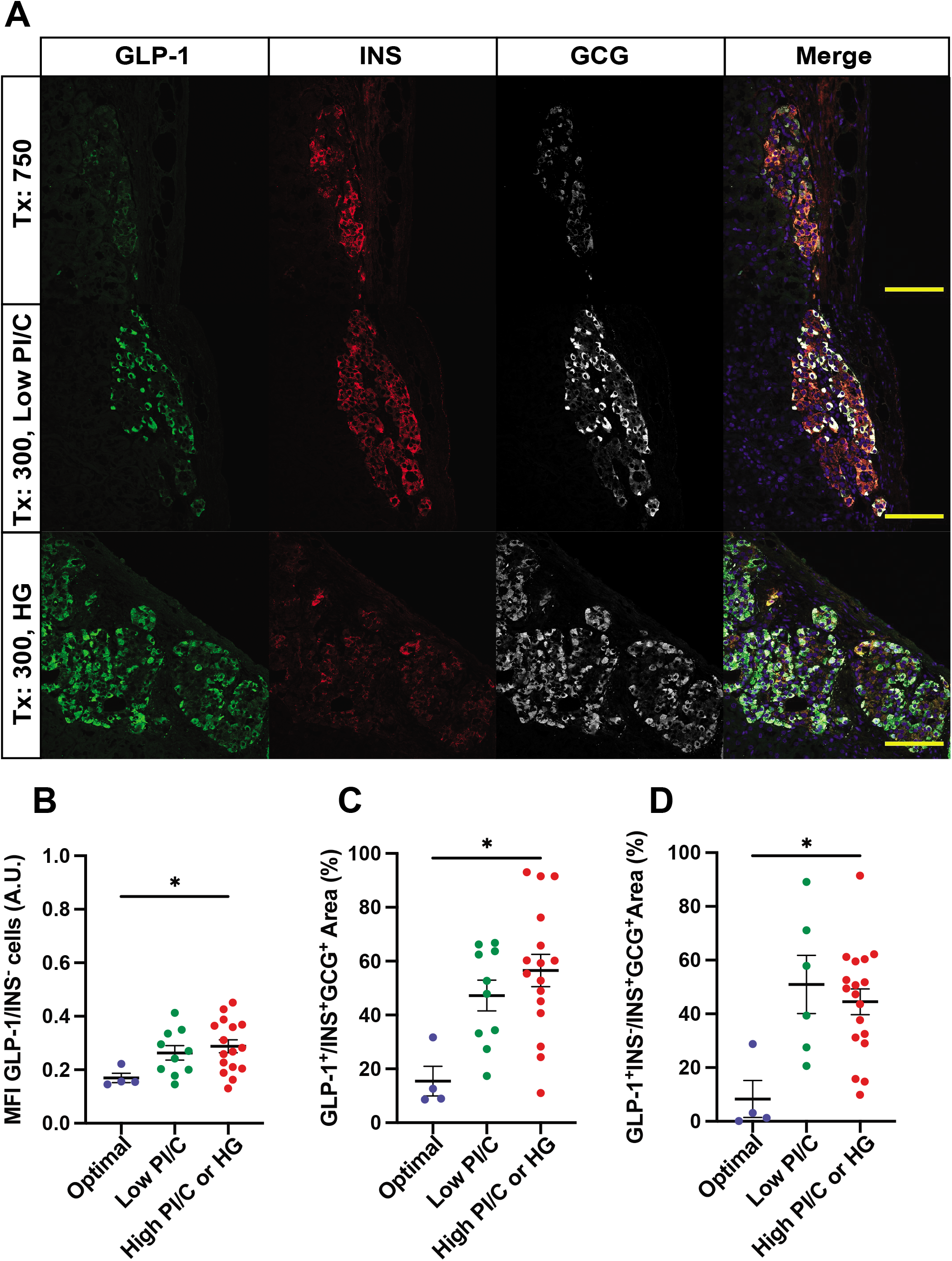
GLP-1-positive area is increased in human islet grafts from mice that displayed high PI/C ratios or hyperglycemia. A) Islet grafts were analyzed by immunofluorescence staining using antibodies against insulin, glucagon, and amidated GLP-1. B) GLP-1 intensity in insulin-negative cells, C) GLP-1-positive area in islet (the combination of insulin- and glucagon-positive area), D) and GLP-1 positive insulin-negative area in islet were analyzed.

## Discussion

Islet transplantation is an effective surgical procedure to establish glycemic control in patients with T1D (allogeneic) or following pancreatectomy (autologous); however, transplant outcomes vary with a large proportion of patients requiring insulin therapy a few years after the surgery^1^. In patients undergoing TPIAT, because islet mass is often impaired by a fibrotic and diseased pancreas, only about 30% of patients successfully wean off insulin after transplant. Here we demonstrate that plasma PI/C and proIAPP/IAPP ratios are good indicators of islet autotransplantation outcomes. Elevated PI/C and proIAPP/IAPP ratios are closely associated with higher insulin usage, MMTT AUCglu, and ACRglu at 3 months post-autotransplant. Most importantly, patients who remained insulin dependent at 1-year post-transplant displayed higher PI/C ratios at 3 months post-autotransplant, suggesting that elevated islet prohormone ratios may indicate an islet mass that is metabolically stressed and insufficient to allow insulin independence.

It has been reported that patients transplanted with higher IEQ are likely to remain insulin-free^20–23^. In this study, we showed that lower PI/C and proIAPP/IAPP ratios are associated with higher islet mass transplanted. Interestingly, one subject who received an ample amount of islets (6,926 IEQ/kg) became insulin-dependent within a year post-transplant. We found that this subject displayed an elevated PI/C ratio at 3 months follow-up. We propose that the synergistic use of IEQ and PI/C (or proIAPP/IAPP) ratios may improve transplant outcome prediction, and additional studies with larger patient cohorts are needed to establish threshold prohormone ratios for clinical use. We also observed a negative correlation of PI/C and age of TPIAT. Because the PI/C ratios were comparable in patients with different pancreatitis etiologies, further investigation is required to delineate why younger patients have higher PI/C post-transplant.

The cause of islet dysfunction and elevated PI/C and proIAPP/IAPP ratios remains largely unknown, but may involve changes in prohormone convertase activity associated with islet secretory and/or ER stress^24^. We transplanted varying doses of human islets into STZ-diabetic, immune-deficient (NSG) mice, and analyzed circulating human islet (pro)hormone levels and prohormone processing machinery in human islet grafts. While human islet transplantation effectively reversed hyperglycemia, mice that received 750 human islets had significantly lower plasma human proinsulin and proIAPP levels, and markedly lower PI/C and proIAPP/IAPP ratios compared to mice that received fewer human islets. Interestingly, fasting C-peptide levels were comparable among all groups. Our data suggest that although prohormone processing is less efficient, enough mature insulin is being made to prevent the recurrence of hyperglycemia in most graft recipients, even those receiving a low dose (<300) of human islets. The cause of elevated prohormone levels is presumably unrelated to uncontrolled granule exocytosis, because mice that received different amounts of human islets showed comparable kinetics of glucose clearance, displaying an IPGTT curve that plateaued 15 minutes post-stimulation. Further, mice lacking prohormone processing enzymes have largely intact glucose-stimulated insulin and proinsulin secretion^25^. It is possible that prohormone biosynthesis is increased in mice transplanted with fewer human islets, therefore secretory granule content (mature hormones and increased prohormones) were being released together into the blood. Elevated prohormone levels, however, could be indicators of islet duress, as mice that later became hyperglycemic already displayed higher plasma PI/C ratios at 3 weeks post-transplant.

Our histological analysis of human islet grafts revealed that the expression of the prohormone convertase PC1/3 in insulin-positive cells was comparable in mice that received a high dose (750) of human islets to those that became hyperglycemic or displayed higher PI/C ratios, suggesting increase PC1/3 expression in an attempt to maintain processing during beta cell secretory stress. While it is possible that degranulated^26^ and insulin-low beta cells have reduced PC1/3 expression, we were unable to assess insulin-low cells in our image analysis, and that may contribute to biased PC1/3 expression profiles. Further investigation with multiplexed histological and spatial transcriptomic analysis using cell-type specific markers will aid the identification of beta cells of different states to explore the heterogeneity of PC1/3 expression. It is also plausible that even though beta cell PC1/3 immunoreactivity was comparable among different transplant groups, beta cell PC1/3 enzyme activity was impaired in lower-dose islet transplants, resulting in impaired prohormone processing.

We also found that the composition of islet grafts was altered, with the proportion of glucagon-positive area increased and the proportion of insulin-positive area reduced in hyperglycemic mice or mice that displayed higher PI/C ratios. PC1/3 expression was also evident in glucagon-positive alpha cells. Interestingly, GLP-1-positive areas in islets, quantified by the sum of insulin- and glucagon-positive areas, were increased in hyperglycemic mice or mice with higher PI/C ratios. The overall percentage GLP-1-positive cell population in human islet grafts is within the range of previously published human islet studies^27,28^, highlighting the potential roles of islet-derived incretins as autocrine or paracrine in islet transplantation. While the precise mechanism leading to GLP-1 production in alpha cells remains to be clarified^29–34^, our observations may provide useful information to improve clinical human islet transplantation outcomes. GLP-1 is known to promote the survival of beta cells^35^ and stimulate insulin secretion^35^. Nevertheless, the effects of GLP-1 on metabolic regulation during or after clinical islet transplant remain elusive. Although short term metabolic improvement was observed^36–38^, administration of GLP-1 or dipeptidyl-peptidase 4 inhibitors failed to sustain long-term insulin independence^39–42^. One report suggested that GLP-1 infusion during glucose-potentiated arginine stimulation leads to increased proinsulin secretion in islet transplant and pancreas transplant patients compared to control subjects^43^. Interestingly, GLP-1 may suppress glucagon secretion^44^, especially in high glucose environments such as intrahepatically-located islet grafts^45^. It is possible that islet-derived GLP-1 inhibits glucagon secretion^46,47^ and contributes to hypoglycemia incidence in intrahepatic islet autotransplant patients. Follow-up studies on intra-islet GLP-1 production and secretion, in relation to glucagon release during human islet transplantation, will provide insights to inform therapeutic intervention.

In summary, we demonstrate that plasma PI/C ratio is a good indicator of islet graft function, and may be useful in predicting TPIAT outcome. Our clinical observation was reproduced in a mouse model of human islet transplantation, in which we also showed that glucagon-positive cells area and amidated GLP-1-positive area were increased in grafts from mice that displayed hyperglycemia or increased PI/C ratios. Our study suggests that islet prohormone ratios may have value in assessing islet graft function and predicting insulin dependency of autologous islet transplants, highlighting the potential uses of prohormone biomarkers in evaluating the function and integrity of islet transplants.

## Supporting information

Supplementary Table 1 human islet information

Supplementary Table 2 Antibody information

## Acknowledgement

We acknowledge the patients and organ donors who contributed to this study. This work is supported by JDRF (grant 1-INO-2019-794-S-B) and CIHR (grant PJT-153156) to C.B.V., and JDRF postdoctoral fellowship #3-PDF-2017-373-A-N to Y.C.C.

## Disclosure

The authors of this manuscript have no conflicts of interest to disclose as described by the American Journal of Transplantation.

## Data availability

The data that supports the findings of this study are available from the corresponding author upon reasonable request.

## Supporting information

Additional supporting information may be found online in the Supporting Information section.

## Abbreviations

ACRglu: Acute C-peptide response to glucose
ACRpot: Glucose-potentiated acute C-peptide response to arginine
AUC: Area under the curve
GLP-1: Glucagon-like peptide 1
HOMA: Homeostasis model assessment
IEQ: Islet equivalent
IPGTT: Intraperitoneal glucose tolerance test
MMTT: Mixed meal tolerance test
NSG: NOD-scid IL2Rg^null^
PC1/3: Prohormone convertase 1/3
PI/C ratio: Proinsulin-to-C-peptide ratio
proIAPP/IAPP ratio: proIAPP-to-total IAPP ratio
STZ: Streptozotocin
T1D: Type 1 diabetes
TPIAT: Total pancreatic islet autologous transplant

## Notes

### Competing Interest Statement

The authors have declared no competing interest.

