## Supplementary Table 1 human islet information for "Elevated islet prohormone ratios as indicators of insulin dependency in islet transplant recipients"

**Supplementary Table 1: Islet donor information**

| Donor | Islet ID | Sex | Age | BMI | HbA1c | Cause of death | Experiment ID |
|---|---|---|---|---|---|---|---|
| 1 | HP-18310-01 | M | 38 | 28 | 5.9% | Head trauma | YCC-14 |
| 2 | HP-18830-01 | F | 53 | 26 | 5.4% | Head trauma | YCC-16 |
| 3 | HP-19324-01 | F | 43 | 23 | 5.4% | Stroke | YCC-17 |
| 4 | HP-19333-01 | M | 38 | 25 | 5.3% | Head trauma | YCC-18 |
| 5 | R353 | M | 69 | 23 | N/A | Neurological death | YCC-19 |
| 6 | HP-20066-01 | F | 60 | 26 | 5.1% | Stroke | YCC-20 |
| 7 | HP-21161-01 | M | 43 | 26 | 5.4% | Head trauma | YCC-21 |
