## Supplementary Table 2 Antibody information for "Elevated islet prohormone ratios as indicators of insulin dependency in islet transplant recipients"

| Antibodies | Source | Catalog# | Dilution |
|---|---|---|---|
| Rabbit anti-PC1/3 | Abcam | ab191452 | 1:250 |
| Guinea pig anti-insulin | Dako | IR002 | 1:3 |
| Mouse anti-proinsulin | DSHB | GS-9A8 | 1:50 |
| Mouse anti-amidated GLP-1 | Abcam | Ab26278 | 1:250 |
| Rabbit anti-glucagon | Abcam | ab133195 | 1:500 |
| Guinea pig anti-glucagon | Linco | N/A (discontinued) | 1:250 |
| Mouse anti-glucagon | Sigma | G2654 | 1:500 |
| Rabbit anti-synaptophysin | Abcam | Ab32127 | 1:250 |
| Goat anti-synaptophysin | R&D Systems | AF5555 | 1:250 |
| Alexa Fluor 488-goat anti-rabbit IgG | Thermo Fisher | A-11008 | 1:250 |
| DyLight 488-goat anti-mouse IgG | Thermo Fisher | 35502 | 1:250 |
| Alexa Fluor 594 anti-guinea pig IgG | Thermo Fisher | A11076 | 1:250 |
| Alexa Fluor plus 594 anti-mouse IgG | Thermo Fisher | A32742 | 1:250 |
| Alexa Fluor plus 647 anti-mouse IgG | Thermo Fisher | A32728 | 1:250 |
| Alexa Fluor plus 647 anti-rabbit IgG | Thermo Fisher | A32733 | 1:250 |
